# Pharmacological Activation of NO-Sensitive Guanylyl Cyclase Ameliorates Obesity-Induced Arterial Stiffness

**DOI:** 10.1101/2025.02.17.638762

**Authors:** Enkhjargal Budbazar, Aylin Balmes, Danielle Elliott, Lisette Peres Tintin, Timo Kopp, Susanne Feil, Robert Feil, Tilman E. Schäffer, Francesca Seta

## Abstract

**Objective:** Arterial stiffness, or loss of elastic compliance in large arteries, is an independent precursor of cardiovascular disease (CVD)^1^ and dementia^2^. Akin to anti-hypertensive and lipid-lowering drugs, arterial de-stiffening therapies could be beneficial at decreasing CVD risk. We previously discovered that enhanced cytoskeletal actin polymerization in vascular smooth muscle cells (VSMCs) contributes to increased arterial stiffness^3^. In aortas and VSMCs, we previously found that decreased NO-sensitive guanylyl cyclase (NO-GC), the NO receptor which synthesizes cGMP, caused downregulation of cGMP-dependent protein kinase I (cGKI) and of its target vasodilator-stimulated phosphoprotein (pVASP^S239^), leading to increased cytoskeletal actin polymerization^3^. In the current study, we tested whether activating NO-GC with an NO-GC activator (cinaciguat) modulates pVASP^S239^ and cytoskeletal actin polymerization in VSMCs, thereby preventing obesity-induced arterial stiffness.

**Approach & Results:** Cinaciguat administration (5 mg/kg) to high fat, high sucrose diet (HFHS)-fed mice, our established model of arterial stiffness^4^, (1) decreased pulse wave velocity, the *in vivo* index of arterial stiffness, without affecting blood pressure, (2) increased aortic pVASP^S239^ levels, and (3) decreased the ratio of filamentous (F) to globular (G) actin, compared to vehicle administration. In cultured VSMCs, cinaciguat (10 μmol/L) increased pVASP^S239^ levels and decreased the F/G actin ratio at baseline and after stimulation with the cytokine tumor necrosis factor α (TNFα), used to mimic the inflammatory milieu of HFHS aortas. These effects were abrogated in aortas and VSMCs from mice with smooth muscle-specific cGKI deletion (cGKI^SMKO^), while being mimicked by a cell-permeable cGMP analog (8-Br-cGMP, 1 μmol/L), which also decreased VSMC stiffness *in vitro*.

**Conclusions:** Collectively, our data strongly support the notion that pharmacological NO-GC activation would be beneficial in decreasing obesity-associated arterial stiffness by decreasing VSMC cytoskeletal actin hyper-polymerization. If translated to humans, NO-GC activators could become a viable approach to clinically treat arterial stiffness, which remains an unmet medical need.

## INTRODUCTION

Cardiovascular disease (CVD) remains the leading cause of death globally, claiming more lives each year than all forms of cancer combined^5^. Arterial stiffness, or loss of elastic compliance of large arteries, which occurs with aging and obesity^6^, is an independent risk factor for CVD^1^. Compelling clinical evidence demonstrated that pulse wave velocity (PWV), the gold standard measure of arterial stiffness, strongly associates with hypertension^7^, heart failure^8^, and dementia^9^. Thus, akin to anti-hypertensive or anti-hyperlipidemic medications, aortic de-stiffening therapies may help prevent cardiovascular events. Cellular and extracellular pathogenic mechanisms of aortic wall stiffening have been identified^10^; yet the translation of these pre-clinical findings into targeted therapies remains elusive.

We previously showed that in a mouse model of dietary obesity, arterial stiffness, measured *in vivo* by PWV, increases after two months of high fat, high sucrose (HFHS) feeding, preceding the development of hypertension^4^, consistent with clinical findings^6^. At the cellular level, we reported that excessive cytoskeletal actin polymerization in vascular smooth muscle cells (VSMCs) is a major contributor to increased arterial stiffness^3,11^. We further discovered that increased cytoskeletal actin polymerization in VSMCs is dependent on the NO-sensitive guanylyl cyclase (NO-GC)/cyclic guanosine monophosphate (cGMP)-dependent protein kinase I (cGKI, aka PKGI)/phosphorylated vasodilator stimulated phosphoprotein (pVASP^S239^) signaling cascade^3^. In this study, we sought to determine whether activating NO-GC with a commercially available NO-GC activator (cinaciguat) modulates pVASP^S239^ and cytoskeletal actin polymerization in VSMCs, thereby preventing obesity-induced arterial stiffness.

## MATERIALS & METHODS

Data and supporting material are available to the research community upon reasonable request.

### Model of diet-induced obesity and drug treatments

Mice with a floxed cGMP-dependent protein kinase I allele (cGKI^fl/fl^)^12^, were obtained from Dai Fukumura (Massachusetts General Hospital, Boston) under an MTA between BU and MGH. Smooth muscle-specific cGKI deletion was achieved by breeding cGKI^fl/fl^ mice with transgenic mice expressing a smooth muscle α-actin promoter-driven tamoxifen-activated CreER^T2^ recombinase^13^. At six weeks of age, cGKI^fl/fl^*/*SMA-CreER^T2+^ mice were fed a diet containing tamoxifen (500 mg/kg, TD.130857, Teckland, Inotiv) or a standard chow for five days, to generate mice with smooth muscle-specific cGKI deletion (cGKI^SMKO^ mice; n=12) and littermate controls (cGKI^ctrl^ mice; n=12), respectively. At two months of age, mice received a normal diet (ND: 4.5% fat, 0% sucrose; n=6) or a high fat, high sucrose diet (HFHS: 35.5% fat (lard), 16.5% sucrose; n=18), and water ad libitum for 6 months. ND and HFHS were custom formulated to contain the same macronutrients, except for fat and sucrose (catalog # D09071702 and D09071703, Research Diets, New Brunswick, NJ, USA). After six months of HFHS, cGKI^ctrl^ (n=9) and cGKI^SMKO^ (n=9) mice were randomized to receive vehicle or cinaciguat (5 mg/kg body weight), daily for 3 weeks by oral gavage. Experimental timeline is illustrated in **Fig. 1A**. To note, this dose of cinaciguat was chosen because, reportedly, not having a significant effect on blood pressure^14^. Mice were housed in the Boston University Medical Campus Animal Facility in temperature- and humidity-controlled rooms, on a 12-hour light/dark cycle, in groups of 4-5 mice per cage. Animal procedures were approved by the institutional animal care and use committee (IACUC) at Boston University. Since our previous study on the role of NO-GC/PKG/pVASP^S239^ signaling pathway on VSMC stiffness was conducted on male mice^3^, the current study used male but not female mice.

**Figure 1.**
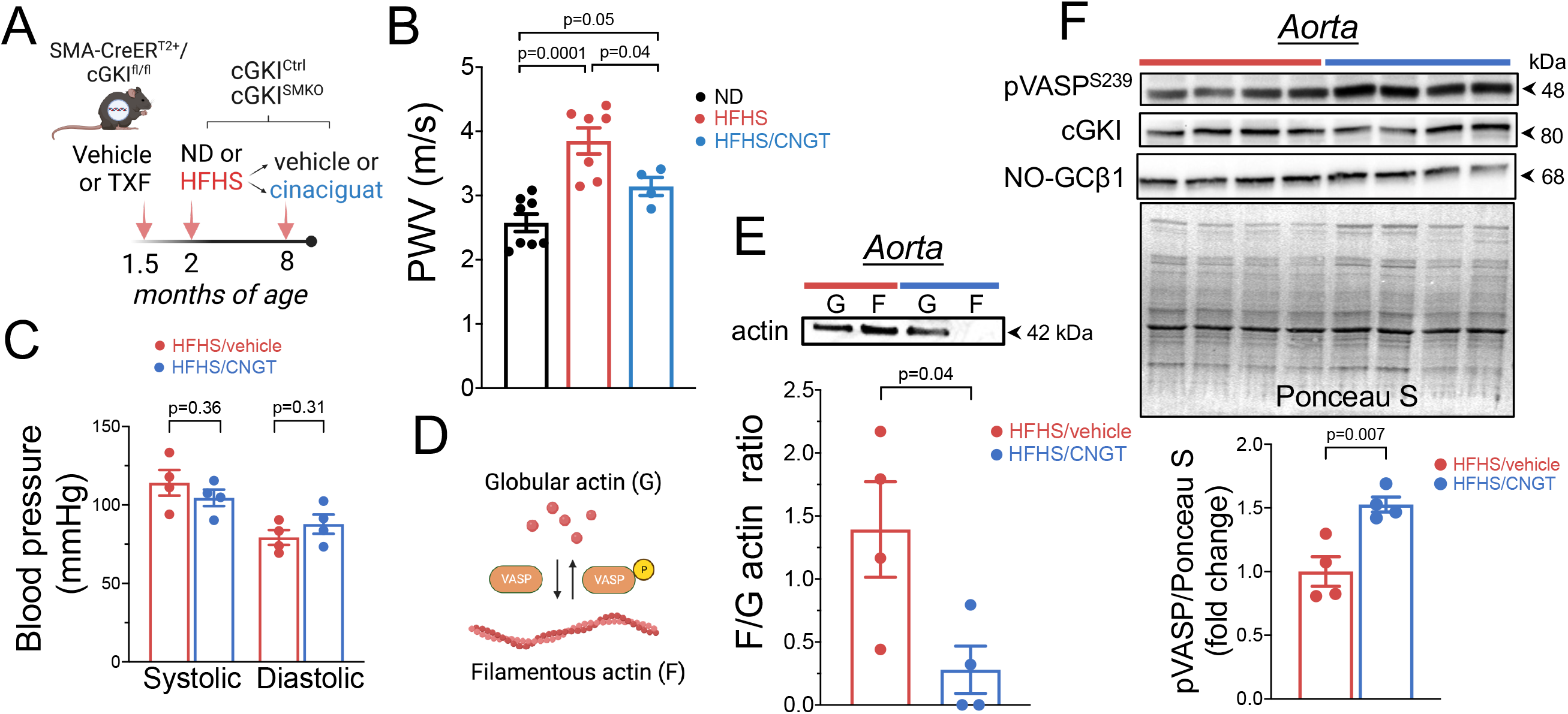
**(A)** Experimental timeline: at six weeks of age, mice with floxed cGKI and smooth muscle α-actin (SMA)-Cre ER^T2^ recombinase (SMA-CreER^T2^/cGKI^fl/fl^) were treated with vehicle or tamoxifen (TXF) to generate mice with smooth muscle-specific cGKI deletion and littermate controls; at 8 weeks of age, mice received normal (ND) or high fat, high sucrose diet (HFHS); after six months, HFHS-fed mice were randomized to receive vehicle or cinaciguat (5mg/kg/d for 3 weeks). **(B)** Cinaciguat administration (5 mg/kg/day) for 3 weeks by oral gavage decreased pulse wave velocity (PWV, m/s), the *in vivo* index of arterial stiffness, in HFHS-fed mice; p=0.04 HFHS vs HFHS/CNGT by one-way ANOVA. **(C)** Cinaciguat administration did not significantly affect systolic and diastolic blood pressure, compared to vehicle; p=0.36 and p=0.31, respectively. **(D)** Filamentous (F) actin is formed by polymerization of globular (G) actin, a process mediated, among others, by VASP and inhibited by phosphorylated VASP. **(E)** Representative Western blot indicates that cinaciguat administration significantly decreased F/G actin ratio in aortas of HFHS-fed mice, compared to vehicle; p=0.04. **(F)** Cinaciguat administration significantly increased pVASP^S239^ levels in aortas of HFHS-fed mice, compared to vehicle; quantitation in graph indicates pVASP^S239^ band intensities normalized to loading control (Ponceau S-stained Western blot membranes) and expressed as fold change of vehicle; p=0.007. Levels of cGKI and NO-GCβ1 in aortic homogenates did not change significantly between vehicle- and cinaciguat-treated HFHS-fed mice.

### Arterial stiffness measurements

Pulse wave velocity (PWV), the *in vivo* index of arterial stiffness, was measured on experimental mice as we previously described^3,4^, at three time points: (1) at baseline (cGKI^Ctrl^, n=12; cGKI^SMKO^, n=12); (2) after 6 months of ND and HFHS (cGKI^Ctrl^/ND, n=3; cGKI^SMKO^/ND, n=3; cGKI^Ctrl^/HFHS, n=9; cGKI^SMKO^/HFHS, n=9); and after randomization of HFHS-fed mice to vehicle or cinaciguat (cGKI^Ctrl^/HFHS/vehicle, n=5; cGKI^SMKO^/HFHS/vehicle, n=5; cGKI^Ctrl^/HFHS/cinaciguat, n=4; cGKI^SMKO^/HFHS/cinaciguat, n=4). Briefly, mice from the various experimental groups were kept recumbent and lightly anesthetized with 1-2% isofluorane on a heated platform to maintain body temperature and heart rate in the 400-500 bpm range during the procedure. A high-resolution Doppler ultrasound (Vevo3100, Fujifilm Visualsonics, Toronto, Canada) was used to image the aorta from the diaphragm to the iliac bifurcation (B-mode, M250 transducer), as we previously described^3,4^. Flow waves and simultaneous electrocardiogram (ECG) were obtained from two locations along the aorta in Power Doppler mode. Acquisitions with unstable heart rate (HR < 400bpm) or unclear QRS peaks in the ECG were excluded from further analysis (4 cGKI^Ctrl^ and 4 cGKI^SMKO^). Arrival times of the flow waves at proximal and distal locations along the aorta were measured by the foot-to-foot method using the R-wave of the ECG as a fiducial point, on at least 5 cardiac cycles for each mouse. PWV was calculated as the ratio of the distance and the difference in arrival times of flow waves at the two locations (m/s).

### Blood pressure measurements

Mice (HFHS/vehicle, n=4 and HFHS/cinaciguat, n=4) were accustomed to the plethysmography system (BP2000, Visitech) by 10-minute training sessions for two consecutive days. On the day of the measurement, mice were carefully handled, to avoid distress, and gently restrained inside dark cassettes placed on a warm platform. Blood pressure tracings were acquired from the middle caudal artery by multiple tail cuff inflation/deflation cycles, for a total of ten minutes for each mouse. Pressure values deviating two standard deviations from the mean were excluded from further analysis. At least twenty systolic and diastolic pressure and heart rate values were obtained for each mouse. Pruned replicates for individual mice were averaged before proceeding to statistical group analysis.

### VSMC culture and cinaciguat treatments

VSMCs were isolated from aortas of 8-week-old cGKI^Ctrl^ (n=3) and cGKI^SMKO^ (n=3) mice by enzymatic digestion with 1 mg/mL collagenase and 2 mg/mL elastase solutions, as we previously described^3^. Freshly isolated VSMCs were seeded on collagen-coated plates and allowed to attach, undisturbed for 3 days, with DMEM containing 1 g/L glucose, 10% fetal bovine serum (S11150, R&D Systems), and 1% antibiotic-antimycotic solution (15240062, 100x, Gibco™). At confluency, VSMCs were sub-cultured in standard culture dishes (Falcon™ 353002, Fisher Scientific) up to passage 5. VSMCs were made quiescent in FBS-free DMEM medium for 24 hours, prior to treatment with vehicle, 10 μmol/L cinaciguat (Sigma-Aldrich) or 1 μmol/L a cell membrane-permeable cGMP analog 8-Br-cGMP (SigmaAldrich) for 15 minutes or 24 hours. In a subset of experiments, VSMCs or aortas dissected from cGKI^Ctrl^ (n=4) and cGKI^SMKO^ (n=4) mice were pre-incubated with 10 μmol/L cinaciguat for 1 hour before stimulation with 10 ng/mL TNFα for additional 24 hours. At the end of the treatment periods, cells and aortas were collected to prepare protein homogenates for Western blot or F/G actin separation, respectively, as described below.

### Western Blot

Aortas and VSMC homogenates or F and G actin fractions were collected in RIPA or LAS2 lysis buffer freshly supplemented with protease inhibitors, and manually homogenized on ice. Protein concentration was measured using the BCA assay (23225, PierceTM BCA Protein Assay Kit, Thermo Fisher Scientific). Equal amounts of protein (25 μg) were separated by SDS-PAGE in NuPAGE 4-12% Bis-Tris gels (NP0335PK2, Invitrogen) or 4-12% Criterion™ XT Bis-Tris (3415023/3415024, Bio-Rad Laboratories, Inc.), then transferred to 0.45-μm pore PVDF membranes. Membranes were incubated overnight at 4ºC with the following primary antibodies: phosphorylated VASP at serine 239 (3114, CST, 56 ng/mL), cGKI (3248, CST, 0.27 μg/mL), sGC isoform β1 (160897, Cayman Chemical, 0.2 μg/mL), pan-actin (AAN01, Cytoskeleton, 0.5 μg/mL), VCAM1 (14694, CST, 0.1 μg/mL), phosphorylated p65-NFκB (3033, CST, 57 ng/mL), GAPDH (2118, CST, 42 ng/mL). The membranes were then washed with TBS-T and exposed to the chemiluminescent substrate ECL (R1002, Kindle Biosciences, LLC) to visualize protein bands using the optical imager iBright (ThermoFisher) with automatic exposure settings. Protein band intensities were analyzed using ImageJ software (NIH, USA) and normalized to GAPDH or Ponceu S-stained membranes, which were used as loading controls. Band intensities from the same experimental group were averaged and the data were expressed in arbitrary units, as fold change of controls, for each replicate Western blot.

### F and G actin separation

Separation of the filamentous (F) and globular (G) actin fractions was performed with G-actin/F-actin *in vivo* assay kit BK037 (Cytoskeleton, USA), following the manufacturer’s recommendations. Briefly, aortas and VSMCs were lysed in LAS2 buffer, then incubated at 37°C for 10 minutes. 100 μL lysates were centrifuged at 350 x *g* for 5 minutes; cleared lysates were then centrifuged at 100,000 x *g* (Optima Max ultracentrifuge, BeckmanCoulter) for 1 hour at 4°C. Supernatants, containing the G actin, were collected in clean tubes while pellets, containing the F actin, were dissolved in 100 μL LAS01 buffer for 1 hour on ice. F and G actin fractions were then flash-frozen in liquid nitrogen and stored at −80°C until Western blot.

### VSMC stiffness

(Young’s modulus) was measured on freshly isolated cells using a commercial atomic force microscopy (AFM) setup (MFP3D-BIO, Asylum Research, Santa Barbara, CA) mounted on an inverted optical microscope (Ti-S, Nikon, Tokyo, Japan) and a cantilever with a pyramidal tip and a nominal spring constant of 10 pN/nm (MLCT-BIO C, Bruker; the spring constant was calibrated using thermal calibration before each measurement). Force maps of 8×8 µm^2^ and 10×10 pixels were recorded centrally on a VSMC with a retract distance of 2 µm, a sampling rate of 2 Hz, and a trigger deflection of 100 nm. Each cell was measured twice (before and after treatment with 8-Br-cGMP, 1 mmol/L, 1 h). Data were analyzed in Igor Pro (WaveMetrics, Lake Oswego, OR, USA).

### Statistical analysis

Statistical analyses were performed with Graphpad Prism v.9.2 software. Datasets were first subjected to a normality test to determine whether they followed a Gaussian distribution. Means of normally distributed datasets were compared using Student’s t-test. Data that did not follow a Gaussian distribution were analyzed using non-parametric tests. Experiments with multiple treatment and genotype groups were analyzed using one-way ANOVA with appropriate post-hoc multiple comparison analysis. *P* values < 0.05 were considered significant. Across-test multiple test correction was not applied. For experiments with VSMCs, replicate experiments utilized individual VSMC preparations from different mice, to ensure biological, rather than technical, replication. Data are expressed as the mean ± SEM and reported as fold-change vs control for each replicate experiment before averaging by genotype or treatment group.

## RESULTS

High fat, high sucrose diet (HFHS) feeding for six months significantly increased arterial stiffness in mice, measured *in vivo* by pulse wave velocity (PWV), compared to normal diet (ND) (2.7 ± 0.1 m/s in ND, n=8 vs 3.9 ± 0.20 m/s in HFHS, n=7, p=0.0001) (**Fig. 1B**), consistent with our previous reports^4,15^. Treatment of HFHS-fed mice with the NO-GC activator cinaciguat (CNGT; 5 mg/kg/day for 3 weeks by oral gavage) significantly decreased PWV (3.9 ± 0.2 m/s in HFHS/vehicle, n=7 vs 3.1 ± 0.1 m/s in HFHS/CNGT, n=4; p=0.04) (**Fig. 1B**). Administration of cinaciguat for three weeks did not induce statistically significant changes in systolic or diastolic blood pressures (SBP: 114.1 ± 8.1 mmHg in HFHS/vehicle, n=4 vs 104.5 ± 5.2 mmHg in HFHS/CNGT, n=4, p=0.36; DBP: 79.3 ± 4.7 mmHg in HFHS/vehicle, n=4 vs 87.8 ± 6.1 mmHg in HFHS/CNGT, n=4, p=0.31) (**Fig. 1C**).

We previously reported that **(1)** aortic stiffness was associated with increased cytoskeletal actin polymerization, measured as filamentous (F) to globular (G) actin ratio, in aortas and VSMCs^3^, and **(2)** F/G actin ratios were inversely correlated with cGMP-dependent protein kinase I (cGKI)-dependent VASP phosphorylation at serine 239^3^ (**Fig. 1D**). Therefore, given that NO-GC activation with cinaciguat should bring about cGKI activation, we assessed whether cinaciguat affects cytoskeletal actin polymerization and pVASP^S239^ levels via cGKI in aortas of HFHS-fed mice. We found that F/G actin ratio was significantly decreased (1.4 ± 0.4 A.U. in HFHS/vehicle, n=4 vs 0.3 ± 0.2 A.U. in HFHS/CNGT, n=4; p=0.04) (**Fig. 1E**), whereas pVASP^S239^ levels were significantly increased (1.0 ± 0.1 A.U. in HFHS/vehicle, n=4 vs 1.5 ± 0.1 A.U. in HFHS/CNGT, n=4; p=0.007) in aortas of HFHS-fed mice after cinaciguat, compared to HFHS-fed mice receiving vehicle (**Fig. 1F**). Cinaciguat administration did not affect the levels of NO-GC (subunit β1) or cGKI (**Fig. 1F**).

Similarly to the aorta, treatment of cultured VSMCs with cinaciguat (10 μmol/L) decreased F/G actin ratio (1.0 ± 0.0 A.U. in vehicle, n=6 vs 0.6 ± 0.0 A.U. in CNGT, n=6; p=0.007) (**Fig. 2A**) and increased pVASP^S239^ levels (1.0 ± 0.0 A.U. in vehicle, n=6 vs 2.9 ± 0.6 A.U. in CNGT, n=10; p=0.009), compared to vehicle (**Fig. 2B**). The effects of cinaciguat on F/G actin and pVASP^S239^ were mimicked by treating VSMCs with a stable cGMP analog (8-Br-cGMP, 1μmol/L) (**Fig. 2A-B**). Likewise, cinaciguat increased pVASP^S239^ in VSMCs treated with TNFα (10 ng/ml), an inflammatory cytokine that we previously showed increases in aortas after HFHS feeding^4,16^. These pVASP^S239^ increases were blunted in VSMCs with cGKI deletion (cGKI^SMKO^) (1.0 ± 0.0 A.U. in cGKI^Ctrl^/TNFα/vehicle, n=3; 9.0 ± 0.7 A.U. in cGKI^Ctrl^/TNFα/CNGT, n=3; 3.9 ± 0.9 A.U. in cGKI^SMKO^/TNFα/CNGT, n=3; p<0.0001 cGKI^Ctrl^/TNFα/vehicle vs cGKI^Ctrl^/TNFα/CNGT; p=0.0022 cGKI^Ctrl^/TNFα/CNGT vs cGKI^SMKO^/TNFα/CNGT) (**Fig. 2C**, quantitation in **2D**). Inflammatory molecules such as VCAM1 and phosphorylated p65-NFκB remained elevated in response to TNFα stimulation, independently of vehicle or cinaciguat treatment (**Fig. 2E**). The effects of cinaciguat on pVASP^S239^ were abrogated by cGKI deletion in cGKI^SMKO^ aortas treated with TNFα *ex vivo*, compared to TNFα-treated cGKI^Ctrl^ aortas (1.0 ± 0.0 A.U. in cGKI^Ctrl^/TNFα/CNGT, n=4 vs 0.6 ± 0.0 A.U. in cGKI^SMKO^/TNFα/CNGT, n=4; p=0.02) (**Fig. 2F**). Moreover, smooth muscle cGKI deletion increased PWV in cGKI^SMKO^ mice, compared to littermate controls (2.6 ± 0.1 m/s in cGKI^Ctrl^, n=8 vs 3.5 ± 0.3 m/s in cGKI^SMKO^, n=8; p=0.01) (**Fig. 2G**). Cinaciguat failed to decrease PWV in HFHS-fed cGKI^SMKO^ mice, compared to vehicle-treated mice (4.3 ± 0.4 m/s in cGKI^Ctrl^/HFHS/vehicle, n=9 vs 4.6 ± 0.7 m/s in cGKI^SMKO^/HFHS/CNGT, n=4; p=0.74) (**Fig. 2H**). Lastly, treatment with 8-Br-cGMP (1 mmol/L) decreased VSMC stiffness on isolated VSMCs, measured by atomic force microscopy (6.7 ± 1.2 kPa in vehicle, n=25 vs 3.8 ± 1.0 kPa in 8-Br-cGMP, n=24; p=0.002) (**Fig. 2I**).

**Figure 2.**
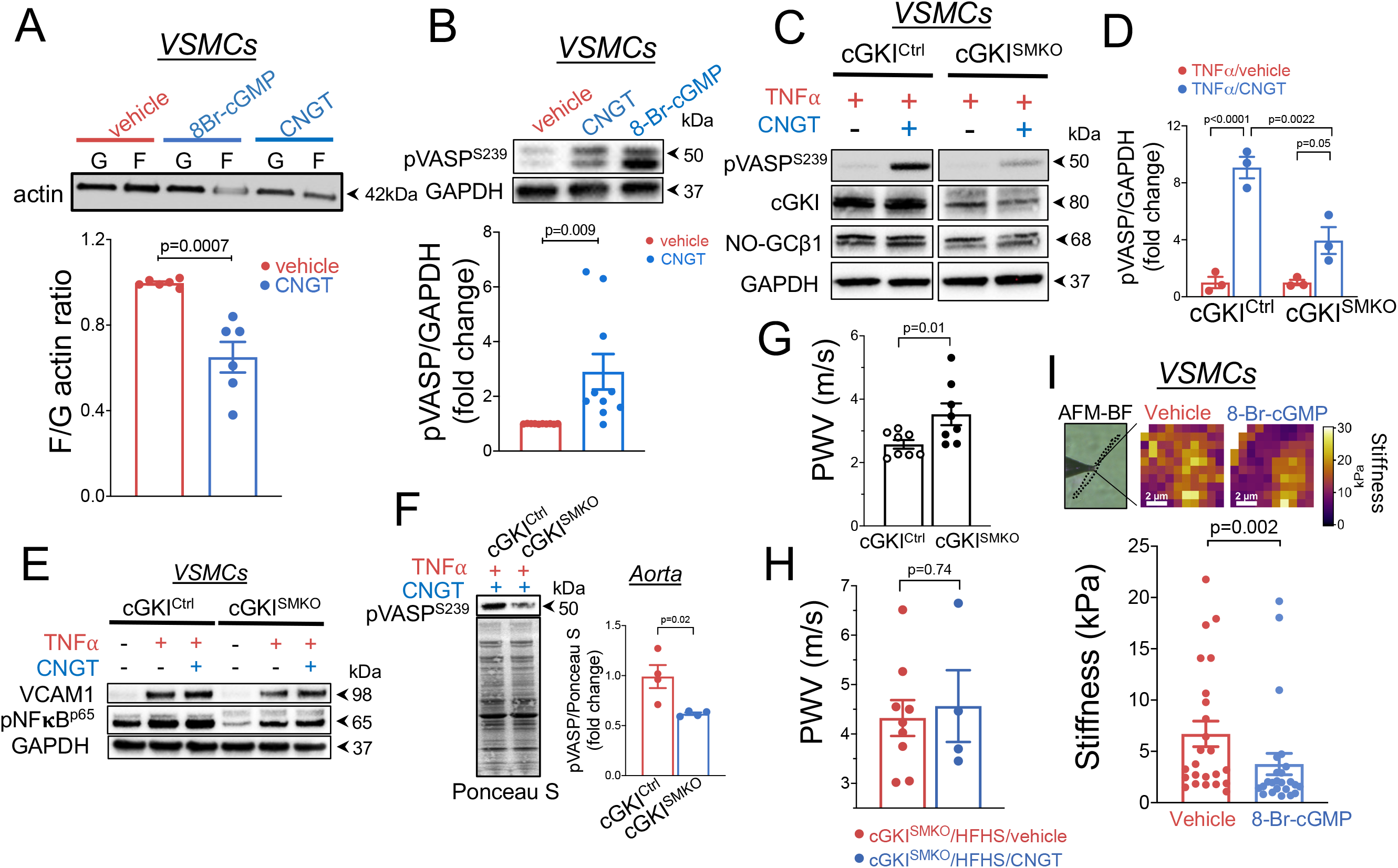
**(A)** Representative Western blot of the F and G actin fractions in VSMCs treated with a cGMP analog (8-Br-cGMP, 1 μmol/L, 1 h) or cinaciguat (CNGT, 10 μmol/L, 1 h); quantitation of band intensities in graph; p=0.0007. **(B)** CNGT (10 μmol/L, 1 h) or 8-Br-cGMP (1μmol/L, 1 h) significantly increased pVASP^S239^ levels in VSMC, compared to vehicle; quantitation in graph indicates pVASP^S239^ band intensities normalized to loading control (GAPDH) and expressed as fold change of vehicle; p=0.009. **(C)** Western blot of pVASP^S239^ levels in VSMCs from control (cGKI^Ctrl^) mice and mice with smooth muscle cGKI-deletion (cGKI^SMKO^) stimulated with the cytokine TNFα (10 ng/ml), or vehicle; quantitation in graph **(D)** indicates pVASP^S239^ band intensities normalized to loading control (GAPDH) and expressed as fold change of vehicle; p<0.0001 cGKI^Ctrl^/TNFα/CNGT vs cGKI^Ctrl^/TNFα/vehicle; p<0.05 cGKI^SMKO^/TNFα/CNGT vs cGKI^SMKO^/TNFα/vehicle; p<0.0022 cGKI^SMKO^/TNFα/CNGT vs cGKI^Ctrl^/TNFα/CNGT. **(E)** CNGT (10 μmol/L) did not significantly affect the expression of inflammatory markers VCAM1 and phosphorylated p65-NFκB in cGKI^Ctrl^ or cGKI^SMKO^ VSMCs stimulated with the cytokine TNFα, compared to vehicle. **(F)** CNGT (10 μmol/L) failed to increase pVASP^S239^ expression in cGKI^SMKO^ aortas stimulated ex vivo with the cytokine TNFα (10 ng/ml), compared to vehicle. **(G)** CNGT (10 μmol/L) failed to decrease PWV (m/s), the *in vivo* index of arterial stiffness, in HFHS-fed cGKISM^KO^ mice, compared to HFHS-fed cGKI^Ctrl^ mice; p=0.74. **(H)** Representative images of atomic force microscopy (AFM) measurement of stiffness (kPa) of VSMCs treated with vehicle or 8-Br-cGMP (1 mmol/L); bright field (BF) image shows an individual cell; heatmap indicates cell stiffness; p=0.002. Scale = 2μm.

## DISCUSSION

When the aorta and other large arteries stiffen with aging or obesity, they lose their intrinsic capacity of buffering the pulsatility of cardiac contraction resulting in poor cardiac perfusion and elevated pulse pressures propagating to the downstream microcirculation, particularly to high flow/low resistance organs, such as the brain and kidney. Structural and functional changes in those organs, known as target organ damage, are a precursor of overt CVD and dementia^17^. Thus, anti-stiffening therapies, akin to anti-hypertensive or lipid-lowering medications, could become clinically relevant to prevent target organ damage, thereby decreasing the incidence of CVD, particularly among increasingly aging and overweight/obese populations.

Here we report that pharmacological activation of NO-GC, the NO receptor in VSMCs, is effective at decreasing obesity-induced arterial stiffness in a mouse model of metabolic syndrome that, we previously showed, closely mimics the human pathology^4^. Our novel findings demonstrate that **(1)** the NO-GC activator cinaciguat decreases aortic stiffness, measured in vivo by PWV, in obese mice; **(2)** cinaciguat increases pVASP^S239^ and decreases cytoskeletal actin polymerization, even in an inflammatory milieu, and **(3)** these effects are dependent on the cGMP-dependent protein kinase I (cGKI, aka PKGI), as demonstrated by using VSMCs and aortas from mice with smooth muscle-specific cGKI deletion (cGKI^SMKO^) (illustrated in the **graphical abstract**). Of note, the aortic de-stiffening effect of cinaciguat was independent of blood pressure because the cinaciguat dose we employed did not significantly affect blood pressure (**Fig. 1C**), consistent with previous reports^14^.

NO-GC is a crucial enzyme for cardiovascular homeostasis^18^. Upon binding of the endogenous ligand NO to its ferrous heme moiety, NO-GC synthesizes cGMP, a second messenger required for VSMC relaxation. In the settings of oxidative stress, when NO bioavailability is diminished and/or the heme moiety is oxidized to its ferric form and unable to bind NO, NO-GC becomes inactive. Hence, NO-GC is an attractive pharmaceutical target for CVD, in which oxidative stress and low NO bioavailability are generally a major culprit. Several NO-GC modulators, classified as NO-GC stimulators and NO-GC activators based on their pharmacodynamic properties, have been developed^19^. NO-GC stimulators, of which riociguat is the lead compound, are able to activate native NO-GC, but not its oxidized/heme-free form^19^. In contrast, cinaciguat and other NO-GC activators are able to selectively activate the oxidized/heme-free enzyme, thereby stimulating cGMP synthesis even under pathological conditions in tissues containing oxidized, NO-insensitive NO-GC^19^. Because of their unique pharmacodynamic property, we posited that, in the vasculature of HFHS-fed obese mice, whose NO bioavailability is drastically decreased and oxidative stress significantly increased^4^, a NO-GC activator would sustain cGMP production more efficiently than a NO-GC stimulator. Hence cinaciguat was the drug of choice in the current study to determine the effects of NO-GC activation on obesity-induced arterial stiffness. We did not test other NO-GC modulators. Therefore, we cannot generalize our findings to other NO-GC activators or stimulators. Moreover, the translational value of cinaciguat is currently limited by its low solubility, poor bioavailability, and short half-life requiring frequent administrations^19^. The development of NO-GC activators like runcaciguat with improved bioavailability compared to cinaciguat^20^, will be crucial to translate our pre-clinical findings to the clinic. Our previous study on the role of NO-GC/cGKI/pVASP^S239^ cascade in VSMC actin polymerization and stiffness was conducted with male mice^3^. Therefore we did not use female mice, a limitation of the current study. Additional studies to investigate potential sex dimorphism in the effects of cinaciguat or other NO-GC modulators on arterial stiffness are essential.

Polymerization of non-muscle cytoskeletal actin ¾mainly β- and g-actin, can increase VSMC stiffness and VSMC tension development in response to mechanical stimuli, thereby sustaining vessel tone and diameter^21^, independently of myosin light chain phosphorylation and actino-myosin cross-bridge cycles^22^. We^3,11^ and others^23^ demonstrated that excessive VSMC cytoskeletal actin polymerization and VSMC stiffness contribute to aortic wall stiffening suggesting regulation of actin polymerization as an appealing, albeit challenging, site for therapeutic intervention^24^, as recently suggested also by others^10,25^. A major mediator of actin filament polymerization, VASP interacts with vinculin, zyxin, and profilins^26,27^, all crucial components of actin filament assembly at the focal adhesions, which are subcellular sites of actin filament attachment to the extracellular matrix, essential for maintaining a mechanically competent aortic wall. VASP overexpression has been shown to induce F-actin assembly^28^ whereas VASP phosphorylation at serine 239 is sufficient to inhibit actin filament polymerization^29^. We previously demonstrated that aortic VASP^ser239^ phosphorylation is decreased in models of arterial stiffness^3,15^. Overall, our findings that cinaciguat decreases F/G actin ratios in aortas and VSMCs, and arterial stiffness *in vivo*, are consistent with cinaciguat increasing levels of pVASP^ser239^, as we similarly reported in platelets^30^. Moreover, our study highlights a crucial role of smooth muscle cGKI in regulating pVASP^ser239^ and arterial stiffness, as demonstrated by the fact that smooth muscle cGKI deletion increases PWV in young cGKI^SMKO^ mice (**Fig. 2G**). Our study also demostrates that cGKI is a downstream effector of cinaciguat since cinaciguat failed to decrease PWV in HFHS-fed cGKI^SMKO^ mice (**Fig. 2H**). Our results do not exclude that cGKI may regulate actin polymerization via additional mechanisms, other than pVASP. For instance, cGKI is able to phosphorylate cofilin1, an actin depolymerizing factor, resulting in cofilin1 inhibition and decreased filamentous (F) actin^31^. Further studies are warranted to explore these cGKI-dependent mechanisms in VSMCs and how they affect VSMC stiffness. Nonetheless, our brief report demonstrates a beneficial effect of NO-GC activation against arterial stiffness and encourages further studies on the therapeutic potential of the NO-GC/cGKI/pVASP signaling pathway to tackle arterial stiffness, which remains an unmet medical need.

## Non-standard Abbreviations and Acronyms

AFM: atomic force microscopy
CNGT: cinaciguat
CVD: cardiovascular disease
CreER^T2^: tamoxifen-activated Cre recombinase
cGMP: cyclic guanosine 3’,5’-monophosphate
8-Br-cGMP: 8-bromo cyclic guanosine 3’,5’-monophosphate
cGKI: cGMP-dependent protein kinase I
cGKI^SMKO^: mice with smooth muscle-specific cGKI deletion
ND: normal diet
NO: nitric oxide
HFHS: high fat, high sucrose diet
pNFκB: phosphorylated nuclear factor κB
pVASP: phosphorylated vasodilator-stimulated phosphoprotein
PWV: pulse wave velocity
NO-GC: NO-sensitive guanylyl cyclase
SMMHC: smooth muscle myosin heavy chain
TNFα: tumor necrosis factor α
VASP: vasodilator-stimulated phosphoprotein
VCAM1: vascular cell adhesion molecule 1
VSMCs: vascular smooth muscle cells

## FUNDING SOURCES

This work was supported by NIH R01 grant HL136311 to FS; BU CTSI Integrated Pilot grant 1UL1TR001430 to FS; a BU CTSI Voucher Award 1UL1TR001430 to EB; by the Deutsche Forschungsgemeinschaft (German Research Foundation) – Projektnummer 335549539 – GRK 2381 RF/SF/TS; by the Reinhard Frank-Stiftung; and by an Add-on Fellowship of the Joachim Herz Foundation to AB.

### ACKNOWLEDGEMENTS

We would like to thank the Boston University Medical Campus Analytical Instrumentation (Drs. Lynn Deng and Matthew Au) and Cellular Imaging Cores (Dr. Michael Kirber) for their expert technical support. Figures 1A, 1D, and the graphical abstract were created with BioRender.com.

## AUTHORS CONTRIBUTIONS

AB, EB, DE, and LPT performed experiments and reviewed the manuscript; TK provided cells for the AFM measurement; SF, RF, and TS provided constructive criticism to the manuscript; FS designed and coordinated the study, performed PWV measurements, analyzed and interpreted the data, and wrote the manuscript.

